# Implications of recursive Bayesian Sensory Inference

**DOI:** 10.64898/2025.12.30.696730

**Authors:** Erik Skjoldan Mortensen, Maud Eline Ottenheijm, Mark Schram Christensen

**Affiliations:** Department of Psychology, University of Copenhagen, Øster Farimagsgade 2A, Copenhagen, 1353, Denmark

**Keywords:** Proprioception, Sensory integration, Bayesian inference, Sensorimotor control, Muscle spindles

## Abstract

During everyday life, we rely on proprioceptive feedback to inform us of the current position and velocity of our limbs. In this article, we introduce a computational model of arm state inference grounded in Bayesian sensory integration, through which we examine the balance between position-based and velocity-based proprioceptive feedback. The model was developed to compare the behaviour of two contrasting proprioceptive configurations: one relying primarily on position signals and another primarily on velocity signals, reflecting the contributions of type II and type Ia muscle spindle afferents. Our simulations show that a simulated Agent with heavy reliance on velocity-based feedback tends to represent arm movement relative to previously estimated positions, such as when the hand is viewed prior to movement initiation. In a simulated reaching task with briefly presented offset visual feedback, such a system produces the classically observed shifted endpoint errors, while an Agent relying mainly on positional feedback is able to correctly update its inferred hand position after visual feedback is removed, thereby limiting this type of biased endpoint error. When biased visual feedback remains continuously available, or when muscle vibration is simulated, the two configurations can produce very similar behaviour. These findings underscore the importance of carefully considering the composition of proprioceptive signals when building computational models and when interpreting human sensorimotor experiments.

## 1 Introduction

During motor control, the brain tracks both the position and movement of our limbs. Even with our eyes closed, we can bring the tips of our index fingers together, a function that depends on sensory signals from proprioceptive receptors in our muscles (Tuthill and Azim 2018). One set of these receptors is most sensitive to the rate of muscle lengthening, providing information about movement, while another set signals the muscle’s current length, giving us a sense of limb position (Proske and Gandevia 2012). While one might imagine a simple state inference system where positional sensors directly provide us with our sense of the current body pose and limb positions, and velocity sensors provide us with a sense of body and limb motion, it has long been recognised that these signals can in principle be utilised in an integrative fashion to augment one another (Wolpert et al. 1995; Todorov and Jordan 2002; Friston 2010). For example, given a current estimate of hand position, then a signal representing its velocity may be used to predict how the hand position should change in the immediate future. However, there has so far been limited discussion and investigation of the extent to which these two feedback channels are used to augment one another and the relative reliance on each. Computational models of proprioceptive state estimation typically assume a particular balance between position and velocity feedback, but this balance is rarely made explicit, and it is seldom tested directly. We therefore ask whether the sense of limb position depends substantially on velocity-based proprioceptive feedback. We address this by comparing two simulated Agents whose proprioceptive position and velocity channels differ in their relative precision, and asking whether they can be told apart in classic sensorimotor paradigms. Our results suggest that they often cannot, which means such paradigms may be less diagnostic of the underlying feedback balance than they appear, while in one setting they separate clearly and point towards a larger role for velocity feedback than is usually assumed.

## 2 Background

### 2.1 Bayesian inference

Moving our hand to a target requires feedback on the arm’s initial state and how it evolves over the course of the reaching movement. When moving, we have multiple senses to guide us; vision may inform us of both the target location and the location of our hand in a visual reference frame, while proprioception allows us to feel our hand and arm through feedback from muscle, skin, and joint sensors. In addition to these feedback signals, our intention to move forms a feed-forward signal that may likewise inform us of how these different sources of sensory feedback are expected to change over time. Yet it is a basic principle of imperfect and noisy biological sensors and neural communication that these different sources of information will at any given time never perfectly match each other, nor the ground truth, which they are set up to represent (Faisal et al. 2008). Instead, such signals are at best simplified and noisy abstractions, which the central nervous system must process in order to infer the pose and movement of the body and its limbs.

This large array of available, and to some extent mismatching, information highlights the need for a method to integrate different sources of information into a coherent state estimate, which can then be used for motor control. Bayesian sensory integration has been proposed as a way to understand this phenomenon (Knill and Pouget 2004; Franklin and Wolpert 2011). A Bayesian state estimator may, over time, learn the expected level of noise present in a given input sensory channel and output motor channel and use this noise estimate to weigh both incoming and outgoing signals by their expected precision. Several studies have demonstrated human behaviour indicative of such precision-weighted sensory integration, when experimenters have manipulated the availability, reliability, and bias of different sources of sensory feedback (Van Beers et al. 1996, 1999; Ernst and Banks 2002; Körding and Wolpert 2004; Chancel et al. 2016a).

By linking such Bayesian sensory integration with a dynamics model, the framework further describes how the perceived state of the arm is influenced not only by the currently incoming sensory feedback but also by the previously perceived state of the arm. The currently perceived state of the arm represents prior knowledge that future sensory data should be used to update, rather than attempt to construct the whole state of the arm “from scratch” using only the latest sensory feedback. For example, if the arm is believed to be stationary, then the currently perceived arm position represents a prior which incoming sensory feedback may refine, weighted by the expected sensory precision; minor modulation in the positional feedback might either represent true movement or merely sensory noise. If, however, the arm is perceived to move in a particular direction, then changes in the positional feedback corresponding to this movement should be more readily believed. The relative precision of different sensory feedback channels would in this framework dictate the relative trust placed on each source of information.

If a control signal to contract the biceps was recently sent, then the dynamic model could be used to predict both how the elbow should begin flexing in response and how downstream incoming sensory feedback would be expected to change based on this action. Several studies have shown how efferent drive affects the perception of limb position, both when afferent feedback is blocked (Gandevia et al. 2006) and available (Laufer et al. 2001; Smith et al. 2009; Erickson and Karduna 2012), demonstrating a role of intended action for state inference and perception.

This model of Bayesian sensory integration forms the basis for state estimation in both Optimal Feedback Control (Wolpert et al. 1995; Todorov and Jordan 2002) and Active Inference (Friston 2010), two theories that have been central to the field of sensorimotor control over the last few decades. While the above description follows the formulation as used in Optimal Feedback Control more closely in terms of the use of efference copy signals, a similar principle applies in the context of Active Inference. In Active Inference, forward models provide sensory predictions of desired states, with actions serving to minimise prediction errors relative to these. While the computational implementations differ, both approaches share the principle of precision-weighted Bayesian integration of state predictions and sensory feedback.

### 2.2 Proprioception

Proprioception is a broad term that encompasses the perception of the body’s state in terms of posture, position, and movement. In particular, proprioception typically refers to the part of perception arising from a collection of diverse peripheral mechanoreceptors situated in the skin, muscles, and joints (Tuthill and Azim 2018). To limit the scope of our introduction to the subject, we will focus here on two of the more well-studied proprioceptors: the type Ia and type II muscle afferents. These are stretch-sensitive muscle spindles found distributed in skeletal muscles, which provide state-dependent feedback to the central nervous system. They integrate directly with the motor control system at the spinal level through various local reflex circuits, providing low-latency feedback, as well as supplying ascending input to the brain to allow for multisensory central state estimation and movement planning (Proske and Gandevia 2012). The type Ia afferent primarily signals the rate of change of muscle length, providing a velocity-based signal (Prochazka and Gorassini 1998; Roll et al. 2004; Albert et al. 2006), while the type II afferent signals muscle length directly (Botterman and Eldred 1982; Roll and Vedel 1982). In practice, there is likely substantial overlap in the responses of each receptor type, such that they may both, to some extent, encode a position- and velocity-dependent signal, and in some instances also acceleration and further time-derivatives in the case of the type Ia afferents (Botterman and Eldred 1982; Macefield and Knellwolf 2018; Blum et al. 2020). They both encode their respective values in their firing frequency, such that type Ia afferents increase their firing frequency proportionally to the stretch-velocity of the parent muscle, while the type II afferents encode the length of the muscle through proportional firing frequency (Roll and Vedel 1982; Botterman and Eldred 1982; Grill and Hallett 1995).

While absolute position sensors, such as the type II afferents, are of clear value for any attempt to estimate the current pose of the body, they also face principled challenges. An absolute state sensor must be able to encode an extensive range of possible states, inherently limiting the resolution of the signal. They are further susceptible to any potential drift of the signal, which must be accounted for to avoid induction of biased estimates; microneurographic studies reveal substantial decay of the firing frequency of type II afferents already within seconds of a ramp-and-hold stretch of a muscle (Roll and Vedel 1982; Edin and Vallbo 1990). Additionally, many factors in the internal environment affect the firing frequency of muscle afferents, including temperature (Mense 1978), pH (Fischer and Schäfer 2005), tissue pressure (Torell and Dimitriou 2024), and arterial pulsation (Birznieks et al. 2012), all of which change to some degree in response to muscular work. These may all pose a challenge for extracting the position- and movement-related signals of interest, making it non-trivial to disambiguate any actual change in posture from sensory noise or drift due to the aforementioned factors. While velocity sensors, such as the type Ia afferent, are not immune to these effects, their phasic response to any change in joint configuration continues to provide a high signal-to-noise ratio signal of when and how the current state is changing.

Nevertheless, it is seen that, if an imposed movement is kept slow enough to fall below the detection threshold of velocity sensors, then much larger changes in position are required for reliable detection, compared to faster movements (Hall and McCloskey 1983; Taylor and McCloskey 1992). This has led to the general interpretation that type II afferents are likely more involved with the estimation of slow and maintained changes in limb position and posture, while type Ia afferents are more central to the control and estimation of faster movements (Banks et al. 2021).

In a Bayesian filter, velocity and positional sensors may be used to augment each other in a reciprocal manner. Velocity sensors, beyond allowing for the direct measurement of movement speed, also provide a source of information regarding how position changes over time.

By combining this with even a noisy position sensor, the absolute position error is prevented from drifting too far, while short-term relative position changes are represented well due to the velocity signal. Several findings support the idea that the central nervous system may be utilising these signals in this manner. For example, viewing the hand prior to reaching towards a target improves endpoint precision (Rossetti et al. 1994; Elliott and Calvert 1990; Vindras et al. 1998), while movements performed from an offset hand position estimate bias the endpoint position estimate by a similar amount (Körding and Wolpert 2004). This might indicate that the perceived arm configuration and hand position are continuously updated in a manner that depends on a previously perceived position; the perception and execution of relative movements continue to function well, even when absolute position errors have been induced with offset visual feedback. Studies of spontaneous proprioceptive drift of movements performed while lacking visual feedback show a similar pattern of absolute state estimation degrading over time,while relative movements remain well-formed (Brown et al. 2003b; Smeets et al. 2006; Patterson et al. 2017).

## 3 Methods

In the following, we will first present a highlevel conceptual introduction to the simulation model. This is followed by the detailed technical implementation, the specifics of the two proprioceptive configurations and the setup of the three simulated study tasks.

### 3.1 Simulation overview

We simulate an Agent using recursive Bayesian Inference to infer the state of a planar 2-joint arm (see Fig. 1) in a continuous loop, as it moves towards a specified target. For our implementation of recursive Bayesian Inference, we use an Unscented Kalman Filter (UKF) (Wan and Van Der Merwe 2000; Julier and Uhlmann 2004). The UKF is a model-based state estimator that continually predicts and updates its point state estimate, with uncertainty in state estimation represented as Gaussians, while also tracking any correlations in uncertainty. To control the arm, we utilise a Linear Quadratic Regulator (LQR) (Athans 1971), a model-based control algorithm that balances the deviation from the desired state with a control cost, with motor control signals corrupted by signal-dependent noise (Harris and Wolpert 1998). This combination of a Kalman filter-based estimator with LQR control and signal-dependent motor noise is closely analogous to the approach used by Todorov and Jordan (2002). With this model, our focus is on exploring the behavioural effects of various induced sensory biases in the context of Bayesian Inference.

**Fig. 1.**
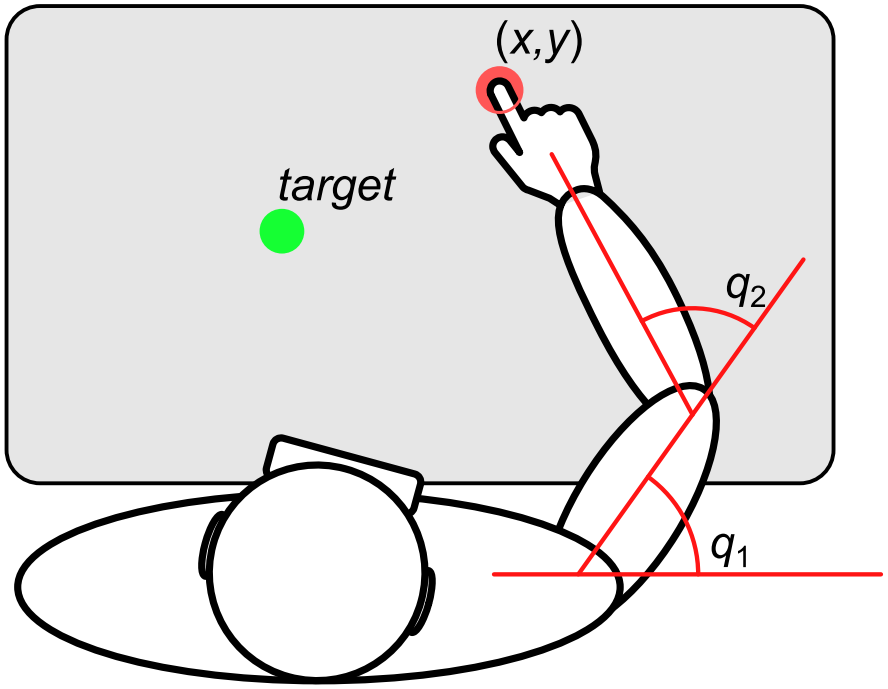
Visualisation of the simulated setup. The Agent is sitting by a table with the weight of its arm supported by air sledges, providing minimal friction to the table surface and allowing only planar movement. It is attempting to infer the state of its arm through noisy proprioceptive and visual sensory feedback. Proprioceptive information is received in the form of the angular positions of the shoulder and elbow (**q** = [*q*_1_, *q*_2_]) and their corresponding angular velocities 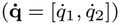. The Agent is outfitted with Virtual Reality glasses, providing visual feedback of only the hand location, which is implemented as a set of 2D Cartesian coordinates, (*x, y*). The Agent is able to produce noisy torques at each joint (***τ*** = [*τ*_1_, *τ*_2_]), in an effort to drive the hand towards a target location

In pseudo-code, the simulation loop is shown in Fig. 2. On each simulated trial, this loop is run for *N* iterations such that 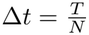, where *T* is the trial duration, with Δ*t* = 10 ms. *k* represents each discretised time step iteration in the simulation.

**Fig. 2.**
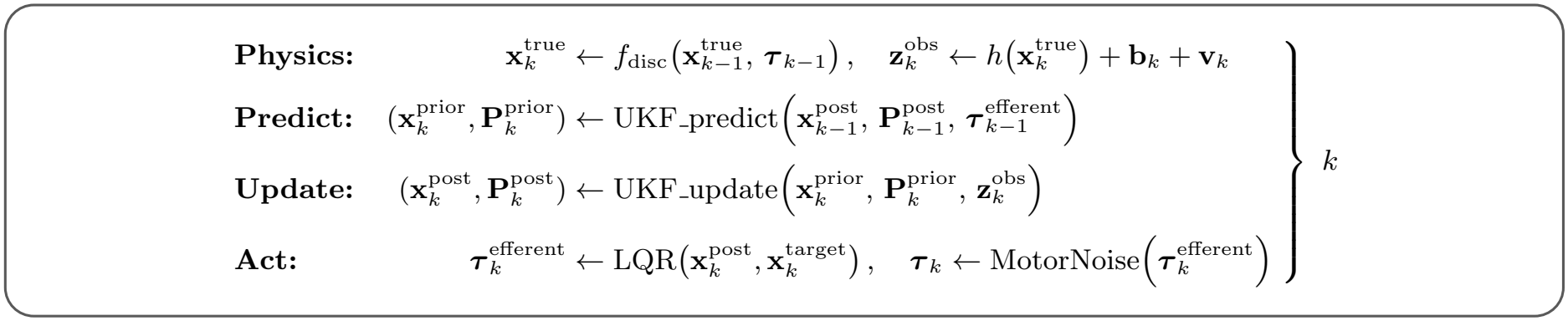
Simplified pseudo-code of the simulation loop. The simulation loop is run for *N* iterations with each step *k* lasting 10 ms. **Physics:** 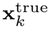 represents the true state of the Agent’s arm at time *t*_*k*_, defined by the shoulder and elbow angular positions and velocities. This state is updated by the discretised physics simulation function *f*_disc_, based on the previous state and muscle-generated torque ***τ*** _*k−*1_. A set of sensory observations 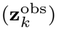 is generated from the updated arm state, based on the state-sensor mapping 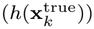 with added Gaussian sensor noise (**v**_*k*_) and possible sensor bias (**b**_*k*_, e.g., a visual offset). **Predict:** The Agent predicts the current state of its arm 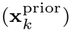 and its uncertainty in this prediction (state covariance, 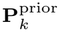) by passing its previous posterior state estimate and uncertainty 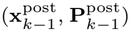, and its associated efferent motor commands 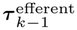 through a dynamic prediction model. **Update:** The Agent updates its posterior state estimate and uncertainty by integrating sensory observations with its predictions. **Act:** The Agent calculates a set of efferent motor commands that will guide its hand towards a specified target state 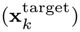. These motor commands are corrupted by signal-dependent motor noise before being applied to the physics simulation on the next step

The Agent continually attempts to estimate the state of its arm and produce motor commands to move its hand towards an indicated target. It lacks direct access to the true state of its arm 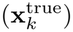, and must instead rely on noisy sensory observations 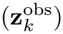. The state of the arm is here fully defined in terms of the shoulder and elbow angular positions and velocities, with the position of the hand further defined by the forward kinematic mapping, given the angular positions and the lengths of the arm segments. The sensory feedback that the Agent receives is of two types: partly proprioceptive feedback, consisting of noisy versions of the angular positions and velocities of each joint, representing a simplified version of the information supplied by type II and type Ia muscle spindle afferents, respectively, and partly visual feedback, here implemented as a set of two-dimensional Cartesian coordinates of the hand position, likewise with added noise.

Together with an internal model of the coupled dynamics of its arm (matching the actual dynamics function *f*_disc_) and its previous efferent motor command, the Agent is able to form predictions about the evolution of the arm configuration (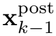to 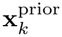), and how certain it is in this prediction 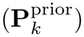. From this predicted state configuration, predictions of expected sensory feedback may be formed, based on both the certainty of the predicted state estimate and the expected level of noise present in each sensor channel; e.g., the Agent may believe visual feedback to be of high precision, and would, in that case, expect the incoming visual feedback to be correspondingly close to the predicted state.

By comparing the expected sensory feedback with the actual sensory feedback 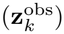, the Agent updates its current state, the posterior 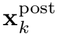 and the associated certainty 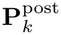.

From this new state estimate, the Agent forms an efferent motor command, here symbolised by 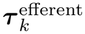, which is corrupted by signal-dependent noise before being applied to the physics simulation of the true arm on the next step.

The simulation was implemented in Python v3.11.5 (Python Software Foundation 2023), using the NumPy v2.3.1 and SciPy v1.13.1 packages. All plotting of the results was done in R v4.5.1 (R Core Team 2024), making use of the tidyverse v2.0.0 (Wickham et al. 2019).

### 3.2 Physics simulation

The arm is set up to move in the horizontal plane and should be considered as supported by a frictionless tabletop, such that gravity is not acting on any joints. The simulated dynamic model of the arm, shown in Fig. 3, is based on the formulations presented in Craig (2005). The state of the arm is defined by **x**^true^ (Eq. P.1), consisting of the angular positions and velocities of the shoulder 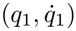 and elbow 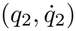 joints. These are related to the position of the hand as described by the forward kinematic mapping **P**_*h*_(**q**) (Eq. P.2). The movement of the arm is simulated as dynamically coupled through the mass term **M**(**q**) and the Coriolis/centrifugal term 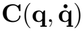, with the generated muscle torques ***τ*** acting on the joints (Eq. P.3–6). Additionally, friction is simulated as viscous damping in the joints with the variable **D**, leading to exponential decay of angular velocity in the absence of applied muscle torques. The dynamics are discretised for simulation using semi-implicit Euler integration (Eq. P.7–8) with a Δ*t* of 10 ms.

**Fig. 3.**
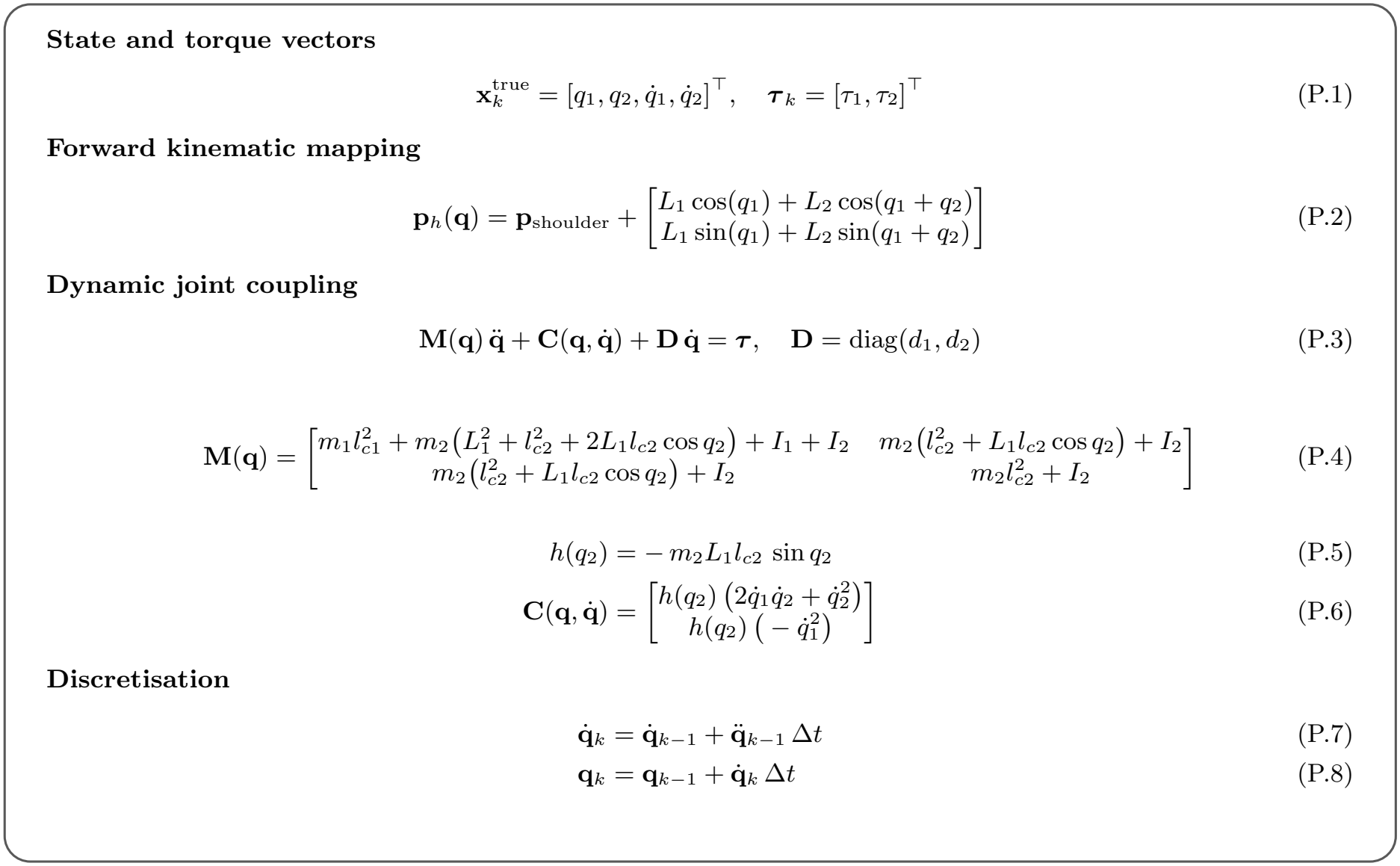
Physics model for the planar two-joint arm. The state comprises the shoulder and elbow angles and velocities; forward kinematics map joint angles to hand position. Joint torques drive coupled dynamics defined by the mass and Coriolis/centrifugal terms, with viscous damping; the dynamics are discretised using semi-implicit Euler integration

### 3.3 Measurement model and motor noise

At each time step *k*, the Agent receives a set of sensory observations 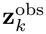, as shown in Fig. 4. These are based on the direct state-sensory mapping, 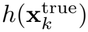, with two added terms (Eq. N.1). Partly, a time-varying bias may be added to simulate the various experimental interventions through **b**_*k*_, e.g., to add a visual offset to the returned visual feedback. Additionally, Gaussian noise is added, given by **v**_*k*_, with the noise of each sensor implemented as time-invariant and independent, given by **R**^true^ (Eqs. N.1 and N.3).

**Fig. 4.**
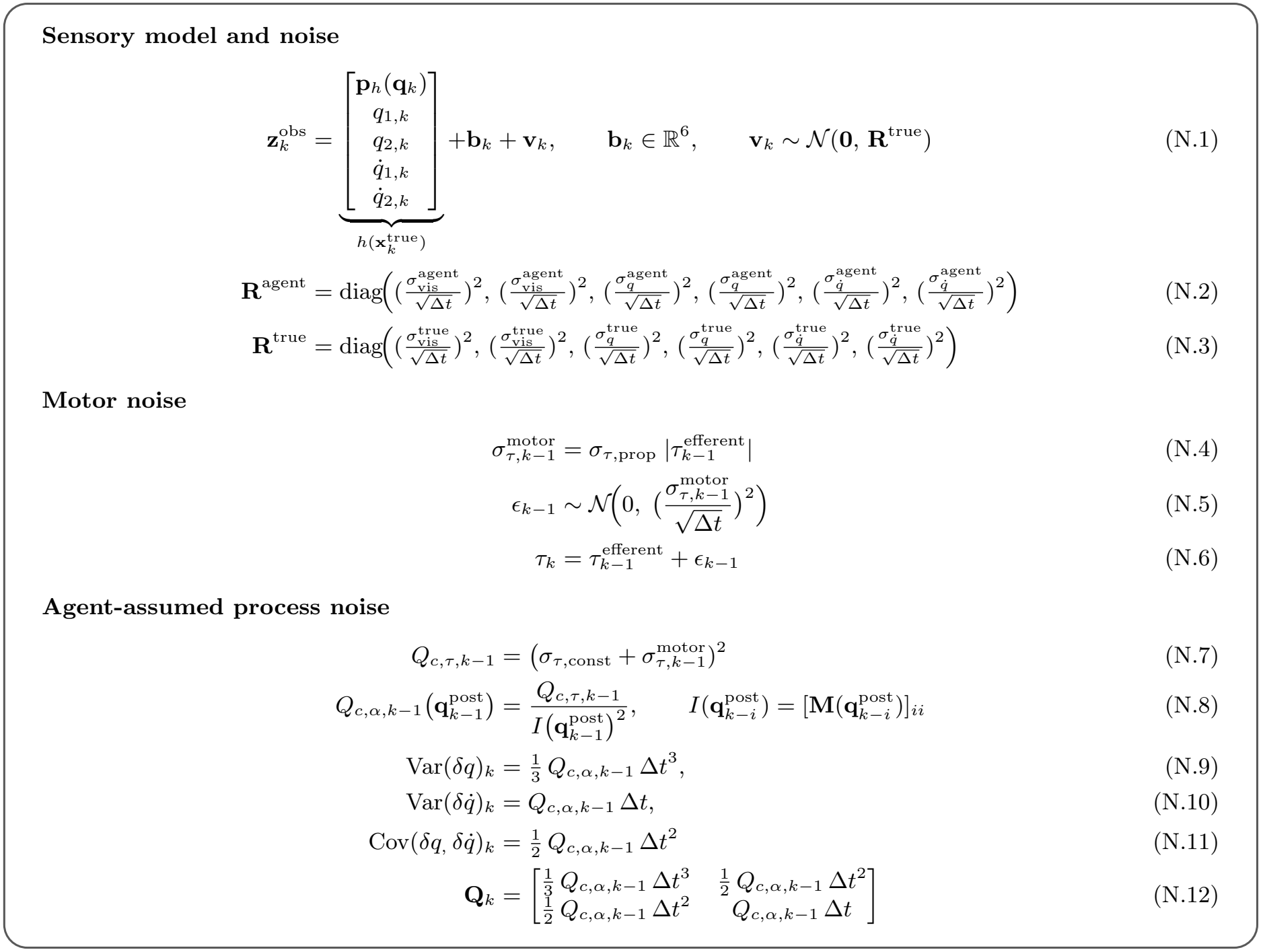
Measurement and noise models. Observations combine hand position and joint angles and velocities with an (optional) additive bias and Gaussian sensory noise. **R**^true^ and **R**^Agent^ denote the true and Agent-assumed sensory covariances (normally equal); noise densities are converted to per-step standard deviations by dividing by 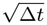. Signal-dependent noise perturbs efferent torque, and the Agent converts its assumed motor and disturbance noise, through joint-acceleration noise, into the discretised process covariance **Q**_*k*_

Motor noise is added to the efferent motor command 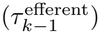 in a signal-dependent manner (Harris and Wolpert 1998), scaling proportionally with the efferent torque (Eq. N.4–6).

Both sensory and motor noise are implemented as continuous-time white noise with a specified power spectral density, which is then discretised for sampling during simulation. This implementation means that noise injection per unit time is independent of the choice of Δ*t*.

During forward prediction to form the prior for the current step, the Agent relies upon the efferent motor command and the previous state posterior estimate. Any uncertainty in the applied torque (Eq. N.7) leads to uncertainty in the angular accelerations (Eq. N.8) applied to each joint, ultimately resulting in increased uncertainty in the angular positions and velocities (Eq. N.12). The Agent-assumed process noise matches the signal-dependent noise, reflecting the true motor noise, with an additional constant term (*σ*_*τ,const*_) meant to represent any unexpected external disturbances. This effectively means it will increase the relative weight of sensory feedback. In a simulation such as this, the actual injected torque noise can be controlled and matched perfectly by the Agent if programmed as such. However, it seems likely that a human performing a motor task should remain somewhat sceptical of their forward predictions from efferent commands or predicted sensory effects of actions to allow for unexpected external perturbations. This is a simple way to implement this effect in a tunable manner, though it should be emphasised that setting the value of this parameter (and other true and Agent-assumed noise parameters) is inherently somewhat arbitrary and can significantly affect the Agent’s behaviour and performance.

### 3.4 State estimation

The Agent estimates its arm state in two steps each loop, using an Unscented Kalman Filter (Wan and Van Der Merwe 2000; Julier and Uhlmann 2004), as shown in Fig. 5. First, it forms a prior by propagating the previous posterior through its internal dynamics model together with the previous efferent motor command (Eq. K.7).

**Fig. 5.**
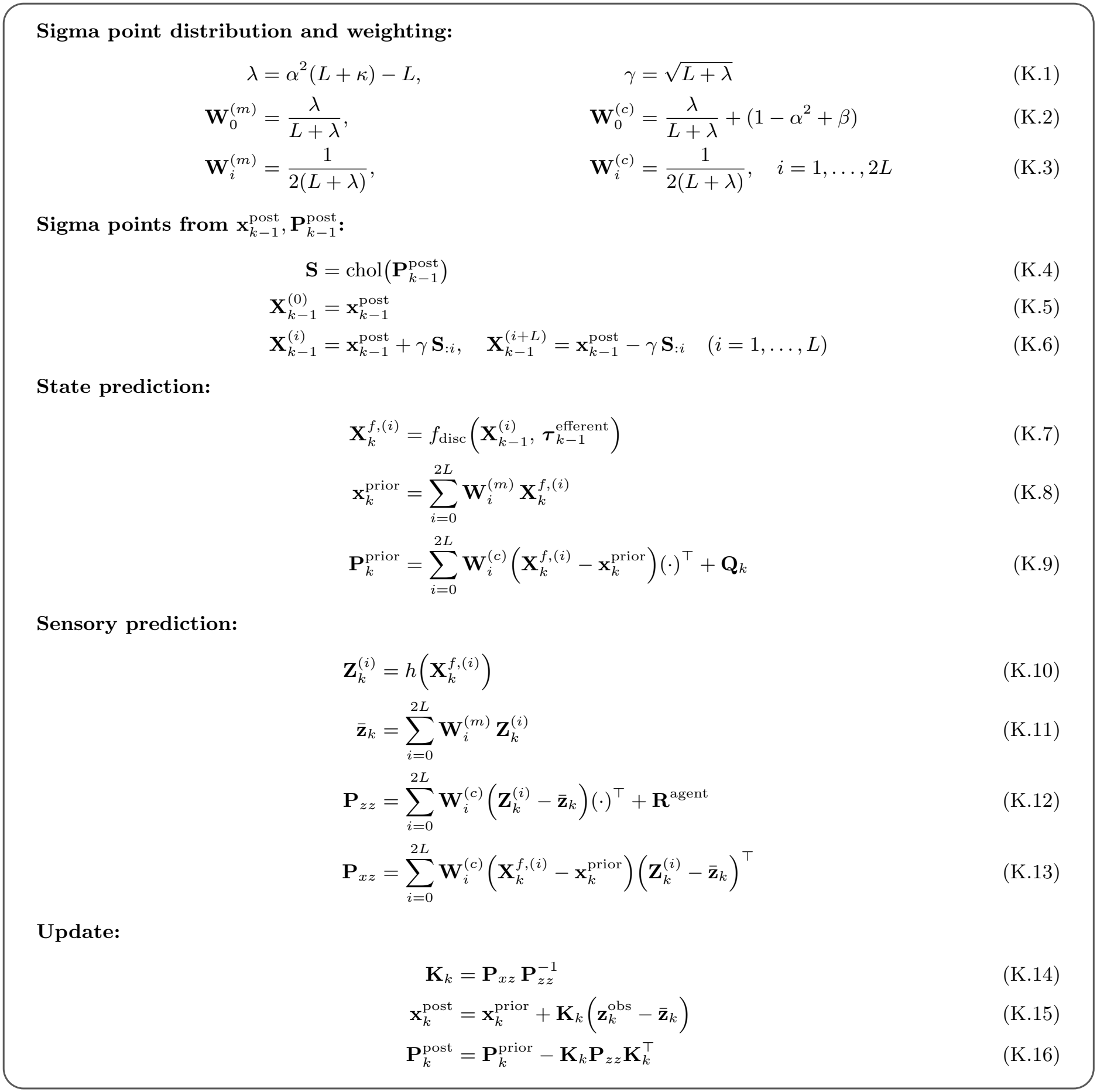
UKF state-estimation procedure. Nine sigma points (*L* = 4) represent the previous state posterior and are propagated through the dynamics and sensory mapping to form state and measurement predictions. The Kalman gain combines the predicted state with the observed measurement to obtain the posterior. The parameters *α, β, κ* determine the sigma-point weights, and *γ* controls their spread

The covariance 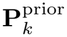 includes the Agent-assumed process noise **Q**_*k*_ (Eq. N.12). 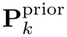 quantifies both marginal uncertainty of each element of 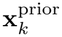 and their expected correlations (e.g., elbow angle and angular velocity may be correlated, so underestimating the angle tends to co-occur with underestimating the velocity during a movement that increases the angle, and so on).

From this prior state estimate, predictions of the expected sensory input are formed (Eq. K.10– 13). These are based on the state-sensory mapping (*h*(**x**)), producing both visual (via the forward kinematic mapping, Eq. P.2) and proprioceptive expected observations, given the expected noise of each sensor, given by **R**^Agent^. Note that the Agentassumed sensory noise (**R**^Agent^) is equal to the true sensory noise **R**^true^, with a notable exception in Simulation 3 as later introduced.

The second main step consists of integrating new sensory observations, 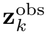. How much new measurements are trusted compared to the predicted state is governed by the Kalman gain **K**_*k*_; when sensory observations are expected to have low noise and correlate well with the true state, then observations are weighted heavily during the update, while high noise sensory feedback will have a smaller impact compared to the prior (Eq. K.14–16).

For a simple 1D example, this corresponds to precision-weighted averaging of the prior and the likelihood (a sensory observation) such that posterior *µ* and *J* (precision, the inverse of variance) would be given by:

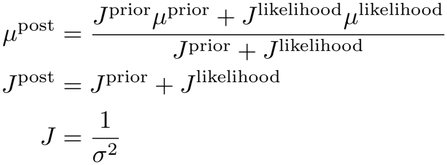

Here *µ*^*likelihood*^ is a sensory identity-measurement, while *J*^*likelihood*^ is the expected precision of this measurement; these would each correspond to 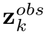 and **R**^*Agent*^.

A key emergent feature of forward prediction is that as long as there is any uncertainty regarding velocity, and how it is changing, then uncertainty about both position and velocity estimates will generally increase, given the process noise assumed here; the Agent will be more unsure about the prior for the next time step than it was about its posterior on the previous step. Conversely, when new information is added in the form of noisy sensory feedback, precision again increases from the prior to the posterior. Across time, the posterior state certainty will approach some equilibrium where the uncertainty gained from the forward prediction is balanced by the information contained in the incoming sensory feedback. This equilibrium will depend on the precision of the available sensory information and the uncertainty associated with the state evolution. For example, in the case of the model here, the precision of proprioceptive positional feedback puts an effective upper limit on how uncertain the Agent can ever be about its position; as seen in the 1D example above, precisions sum, such that the posterior will always have a greater precision than either the likelihood or the prior.

### 3.5 Motor control

Once the Agent has estimated its current state, it computes joint torques to drive the hand toward a target using a Linear Quadratic Regulator (LQR) (Athans 1971), as shown in Fig. 6. At each step, we locally linearise the arm about the current estimate 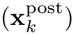 to obtain a discrete model (Eq. L.1–4) and minimise a quadratic cost (Eq. L.5). The finite-horizon problem is solved via the discrete-time backward Riccati recursion to produce feedback gains 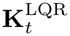 (Eq. L.6–7). We then return the first torque 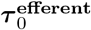 (Eq. L.9), enforcing per-joint rate of force development (Eq. L.10) and torque limits (Eq. L.11). This is repeated at each step in a simplified receding-horizon LQR approximation, which serves as a practical trajectory generator, and optimality for the nonlinear dynamics is not assumed.

**Fig. 6.**
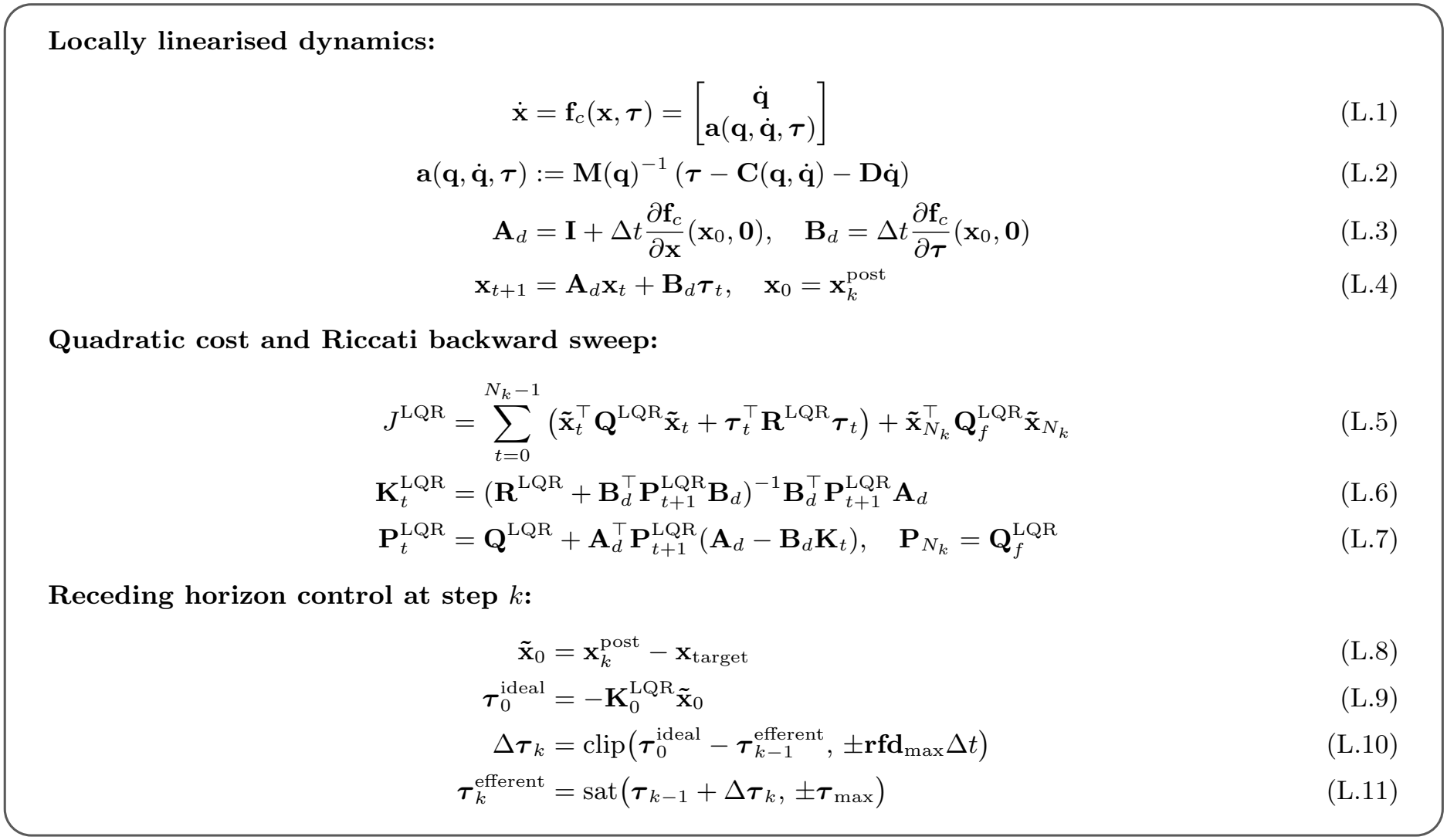
Motor control. The Agent controls its arm using a Linear Quadratic Regulator. At each time step *k*, we linearise the nonlinear dynamics around the current state estimate to obtain **A**_*d*_ and **B**_*d*_ (Eq. L.1–4), the discretised partial derivatives with respect to state and torque changes, respectively. Then the finite-horizon LQR is solved via the discrete-time backward Riccati recursion for *N*_*k*_ steps to obtain feedback gains 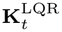 (Eq. L.5–7). *N*_*k*_ is the number of remaining steps at step *k*. We apply only the first computed torque in a receding-horizon manner (Eq. L.9). The weights 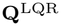 and **R**^LQR^ are fixed, except for the final step, where the terminal weight 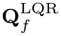 increases the state penalty. Torques are constrained by rate of force development limits, scaled by the time step (Eq. L.10), and a hard saturation (Eq. L.11)

To guide the intended movement time, we specify a time-to-target *T*_*plan*_ such that the planned horizon is 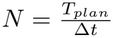, with *T*_*plan*_ reduced by Δ*t* by each step, giving the remaining steps *N*_*k*_ at step *k*. The Agent then attempts to reach the target state (**x**^*target*^; target pose and zero velocity) through an inflated final state deviation weight 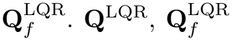, and 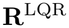are all tunable cost parameters (state and motor, respectively) that will shape the motor behaviour of the Agent. Note that those three parameters, as well as **K**^LQR^, are not related to the previously mentioned process noise (**Q**_*k*_), sensory noise (**R**^true*/*Agent^) or Kalman gain **K**; they are reused here to match standard LQR notation.

### 3.6 Model parameters and simulated tasks

As described in the preceding sections, the Agent relies on several parameters, the configuration of which affects various aspects of the physics simulation, state inference, and motor control. We will here provide an overview of those parameters which have some physical/biological interpretability, with all parameters defined in Appendix table B1. For the physical setup, the length of the upper arm was set to 0.33 m (*L*_1_), and the length of the lower arm (and hand) to 0.45 m (*L*_2_), with their respective masses set to 2 kg (*m*_1_) and 1.7 kg (*m*_2_). Both segments were modelled as uniform rods, with *l*_*ci*_ = *L*_*i*_*/*2 and *I*_*i*_ = *m*_*i*_*L*^2^*/*12, where *I*_*i*_ is the moment of inertia about the segment’s centre of mass, perpendicular to the plane of motion. The shoulder joint is limited to movement between 30^*°*^ horizontal extension and 135^*°*^ flexion, while the elbow is limited to 0^*°*^ extension and 170^*°*^ flexion. For simplicity, both joints experience the same viscous damping of 1.5 N m s, which results in approximate angular velocity half-times of 0.25 s and 0.05 s respectively for the shoulder and elbow joints, meaning that the arm would very quickly slow down in the absence of applied joint torque (shoulder half-time is posture-dependent). These length, mass, and friction values all represent parameters that a human also would likely need to estimate and update as appropriate; for our simulations, we will assume that they are all constant and that the Agent knows and uses the correct values. The time step Δ*t* is set to 10 ms for all simulations.

Visual noise was set to 1 mm s^*−*1*/*2^ per planar dimension, leading to per step sampling standard deviation of 10 mm. Proprioceptive sensory noise is implemented with identical noise densities for both joints. We test and compare two configurations of proprioceptive feedback using the same two noise-density values, assigned oppositely to the position and velocity channels: one configuration relies mainly on velocity-based feedback, with relatively high-precision velocity feedback and low-precision position feedback (high-velocity-low-position, HVLP), and one that relies mainly on position-based feedback, with relatively low-precision velocity feedback and high-precision position feedback (low-velocity-high-position, LVHP). The HVLP configuration has a velocity noise of 0.015 rad s^*−*3*/*2^ and a position noise of 0.06 rad s^*−*1*/*2^. To transform these noise densities for per-step sampling, they are divided by 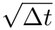; e.g. the above definition indicates that velocity noise here would have a standard deviation of 0.15 rad s^*−*1^ per 10 ms step. The LVHP configuration has the two noise densities flipped, such that it has a velocity noise of 0.06 rad s^*−*3*/*2^ and a position noise of 0.015 rad s^*−*1*/*2^. These values were chosen to highlight both similarities and differences of velocity-dominated and position-dominated configurations of proprioceptive feedback in the chosen simulated tasks, while keeping the performance in terms of endpoint accuracy and precision similar in the non-perturbed control conditions. As such, visual feedback is assumed have fairly high effective precision compared to proprioceptive positional precision for both configurations.

The signal-dependent motor noise scalar is set to 0.05 s^*−*1*/*2^, which leads to a standard deviation of the actual torque corresponding to 5 % of the efferent torque across a 1 second interval (per-step sampling standard deviation of 50 %, similar to the values used by Todorov and Jordan (2002)).

Efferent torque values were limited by both max torque values (50 N m for the shoulder and 30 N m for the elbow), as well as rate of force development limits such that a minimum of 250 ms would be required to reach max torque (max Δ torque of 2 N m and 1.2 N m for the shoulder and elbow respectively per time step).

All simulations were initialised with the Agent perceived state matching the true state 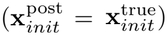, and its state covariance matrix 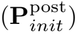 set with sigmas of 1.0^*°*^ and 1.0 ^*°*^ s^*−*1^ respectively for position and velocity uncertainty for both joints (all non-diagonals initialized to 0).

#### 3.6.1 Simulation 1

For Simulation 1, we simulate how the HVLP and LVHP proprioceptive configurations each respond to antagonist vibration during movement. We conceptually match the experiment described by Cody et al. (1990), although we simulate elbow rather than wrist movements. Their results, along with a range of related studies (e.g. Goodwin et al. 1972; Eklund 1972; Roll and Vedel 1982; Gilhodes et al. 1986; Inglis and Frank 1990) demonstrated the classically observed pattern of biased endpoint errors and reduced movement speed in response to various durations and amplitudes of antagonist muscle vibration during active movement, when visual feedback of the hand is withheld. Through varying vibration amplitude, they found that increasing intensity caused successively increased undershooting of the target and larger decreases in movement speed. Varying the point at which the vibration motor was turned on also affected the degree of undershooting; starting vibration earlier resulted in greater undershooting for a given intensity. It is generally recognised that muscle vibration stimulates both type II and type Ia muscle spindle afferents to some extend (Burke et al. 1976), potentially biasing both sensory channels. The perception of the limb is biased such that it corresponds to the vibrated muscle being stretched out further than it is, perceived as a positional and/or velocity offset.

In the simulated setup, the shoulder joint is locked in place, with a target requiring 30^*°*^ of elbow extension to reach it. The allowed movement time is set to 1 s, matching the setup presented by Cody et al. (1990). For the HVLP Agent, we test simulated muscle vibration implemented as a bias of sampled velocity, and for the LVHP Agent, we test it implemented as a position sampling bias. Biasing the non-dominant feedback type would have negligible effects, so we chose to split them up as such to highlight how each of the two configurations would each be affected.

The simulated vibration is implemented through the relevant elbow position and velocity offset being applied to **b**_*k*_ (see Eq. N.1). We test low, medium, and high vibration intensities for each of the two implementations: a positional offset of -5 °, -10 °, or -15 °, and a velocity offset of -5 °/s, -10 °/s, or -15 °/s, respectively. Additionally, we test the medium vibration intensity with onset at 0 °, 10 °, and 20 ° into the movement. The direction of the applied bias is set to match the effect of antagonist vibration, such that it produces a signal that implies greater-than-actual elbow extension/extension velocity.

#### 3.6.2 Simulation 2

In Simulation 2, we simulate how the two Agent configurations each behave when visual feedback of the hand is available throughout the movement, following the setup by Fourneret and Jeannerod (1998). In that study, participants were tasked with making forward reaches towards a target at a distance of 22.2 cm with continuous cursor feedback, corresponding to hand position; upon passing the 5.7 cm mark, a slight visual rotation of 0–10 ° to either side was applied to the cursor. This caused corrections to the reaching movements in the opposite direction, such that the cursor was correctly guided to the target location; after each trial, particiants were asked to report their actual movement direction by choosing from an array of arrows. Here, the majority of participants indicated the direction that corresponded to the observed rotated visual feedback, rather than the direction their hand travelled as they compensated for the incorrect visual feedback, and the reported movement direction was always less than the actual induced offset. Our goal with this simulation was to expand on their findings by considering the posterior of the proposed Bayesian process as what might be available for conscious report, rather than the raw data originating from either visual, proprioceptive, or efferent motor command sources.

We match the experimental setup by applying a visual rotation after the hand passes 5.7 cm, which is implemented through an offset to **b**_*k*_ dependent on the hand’s position. The induced visual offset is gradually applied, with a final off-set of approximately 2.9 cm for a 10^*°*^ rotation once the target is reached. Trials are simulated with a target duration of 2.5 s, which approximately matches the reach durations reported by Fourneret and Jeannerod (1998).

#### 3.6.3 Simulation 3

In Simulation 3 simulated a reaching movement with briefly presented visual feedback partway through the movement, following the setup presented by Körding and Wolpert (2004).

Participants in the experiment was required to reach towards a target positioned 20 cm in front of their start hand position. Visual feedback of the hand position was withheld during the movement, except for 100 ms following the hand passing the halfway (10 cm) mark, forcing them to rely on proprioceptive feedback and efferent output for state estimation for most of the movement. When visual feedback was presented, it could be offset either 0, 0.5, or 1 cm laterally to the left or right, and might either have no, medium, or high blur added to it. They demonstrated that participants would correct their movement so that the implied cursor, rather than their true hand position, was guided to the target, despite the brief period of visual feedback. When visual feedback was blurred, movement adjustments were reduced, indicating a still-present but less influential integration of blurred visual feedback, inline with Bayesian sensory integration assumptions Körding and Wolpert (2004).

Our goal with this simulation was to compare how the precision of brief visual feedback affects the posterior state estimate for each of the two Agent configurations. Visual blur is implemented through increasing Agent-assumed visual noise in **R**^Agent^; this effectively means that the Agent treats blurred visual observations as less informative, decreasing the visual Kalman gain and therefore relying relatively more on its state prior (and through it efferent motor commands) and proprioceptive feedback (see Eq. K.12–16). Specifically, we test for a “No blur” condition where 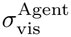 matches the true visual noise of 1.0 mm s^*−*1*/*2^, a “Medium blur” condition where 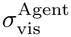 is increased to 1.5 mm s^*−*1*/*2^, and a “Large blur” condition where it is increased to 3.0 mm s^*−*1*/*2^.

We simulate five different lateral visual conditions of no offset, or 0.5 cm or 1 cm to either side; these are applied through **b**_*k*_, and our focus is on comparing the HVLP and LVHP configurations to the original pattern of results. Simulations are here made with a movement duration of 2 s; while Körding and Wolpert do not report the movement duration characteristics, a more recent replication used a maximum allowed time of 5.0 s (Hewitson et al. 2018).

## 4 Results

### 4.1 Simulation 1: Single-joint reaching task with antagonist vibration

Fig. 7 shows the averaged elbow angular position plotted across a 1 s movement, for each of the described Agent configurations. These are split across two possible implementations of simulated muscle vibration as either a bias of the sampled angular position (Fig. 7**b** and **d**) or velocity (Fig. 7**a** and **c**). Here, we aim to highlight that both tested implementations of simulated muscle vibration can produce very similar movement patterns, depending on the assumed relative reliance on position- and velocity-based proprioceptive cues. While the results here are not fully identical, either would be a good match to the observations made by Cody et al. (1990); whether vibration is implemented as a velocity-bias for the mainly velocity-dependent HVLP Agent configuration or as a position-bias for the mainly position-dependent LVHP configuration, both produce the same pattern of undershooting the target.

**Fig. 7.**
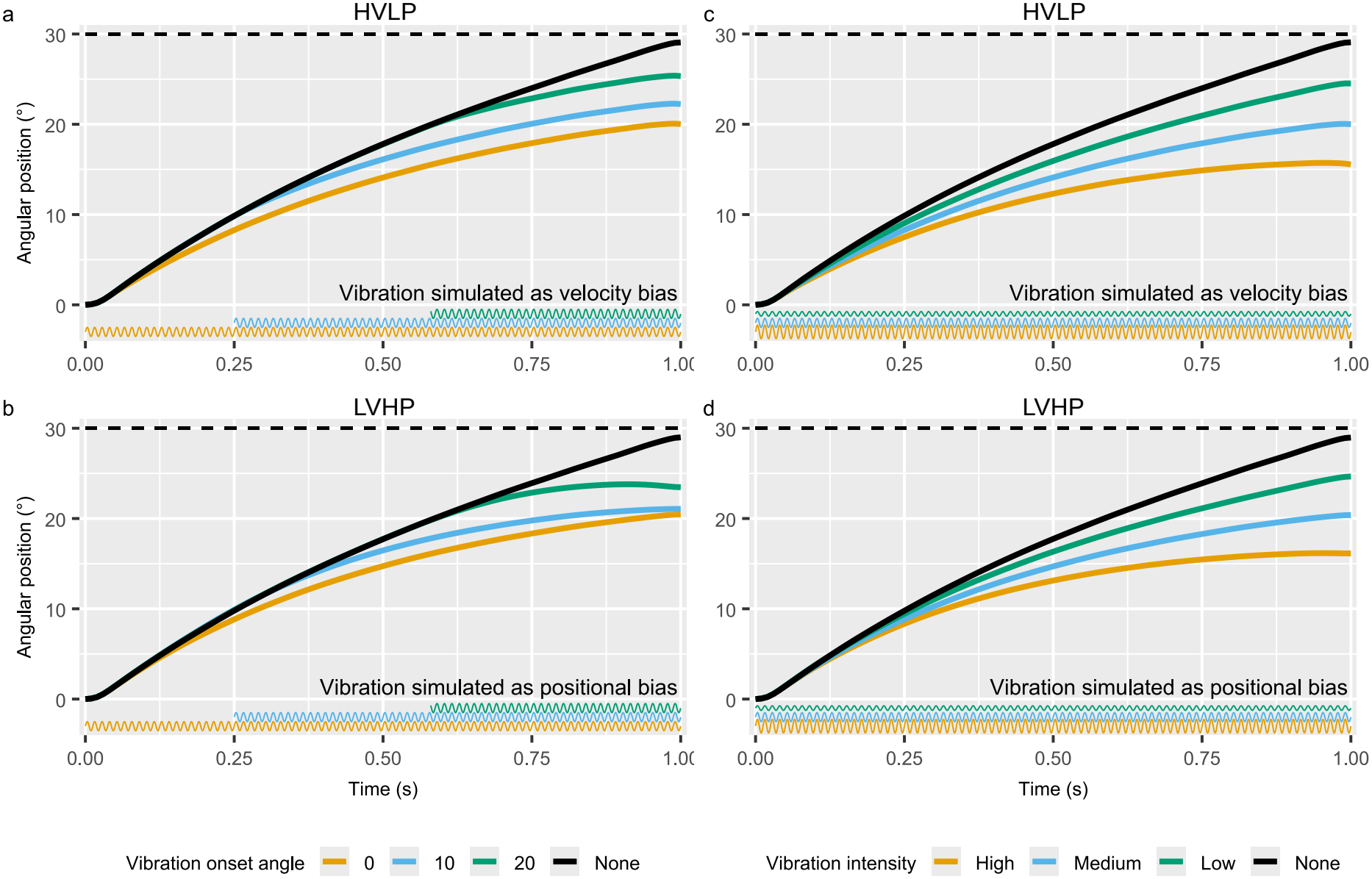
Results of simulation 1. Shows the angular position over a 1-second elbow extension movement towards a target 30° away, analogous to the setup presented by Cody et al. (1990). 0^*°*^ refers to the start position, while the dashed line at 30^*°*^ is the target position. Panels **a** and **b** show the effect of simulated antagonist vibration onset partway through the movement, as illustrated by the corresponding coloured sinusoidal traces along the bottom of each plot. **c** and **d** show the effect of varying the simulated vibration intensity across the whole movement, likewise illustrated by the amplitude of the sinusoidal traces. **a** and **c** are simulated with the HVLP configuration, with simulated vibration implemented as a velocity bias. **b** and **d** are simulated with LVHP configuration, and with simulated vibration implemented as a position bias. The vibration intensity used in **a** and **b** matches the Medium vibration intensity in **c** and **d**. Plotted paths represent a mean of simulated 500 trials each. See the Methods section for details of simulation

The baseline performance of the two Agent configurations is closely matched; the HVLP configuration had a mean endpoint of 29.0^*°*^ (SD = 0.76), while the LVHP configuration had a mean endpoint of 29.1^*°*^ (SD = 0.86), in the nonvibrated condition. In both configurations and all vibration conditions, the Agent correctly guides its posterior perceived hand to the target position, i.e., from the point of view of the Agent, it reaches the target (see Fig. **??**). It may also be noted that if the simulated vibration bias were applied to the non-dominant source of proprioceptive information, e.g., a position bias to the HVLP configuration, little to no effect would be observed, as one might expect. We chose to report the here-presented combinations to highlight how quite different configurations of proprioceptive precision and simulated vibration can produce very similar motor behaviour.

### 4.2 Simulation 2: Reaching task with rotated visual feedback

In Fig. 8, we plot the averaged path of the visual cursor (green), the Agent-perceived posterior path (orange), and the true path of the hand (blue) as the Agent reaches towards the target, for three different visual clockwise rotations of 0^*°*^, 5^*°*^ and 10^*°*^. We also plot the averaged path during the 0^*°*^ condition, as a point of comparison for the remaining conditions (dark grey). While Fourneret and Jeannerod (1998) instructed their participants to follow a straight-line path, we did not implement an explicit straight-line-preference function in our motor control function; the slightly curved path seen in Fig. 8 emerges from the implemented LQR motor control function.

**Fig. 8.**
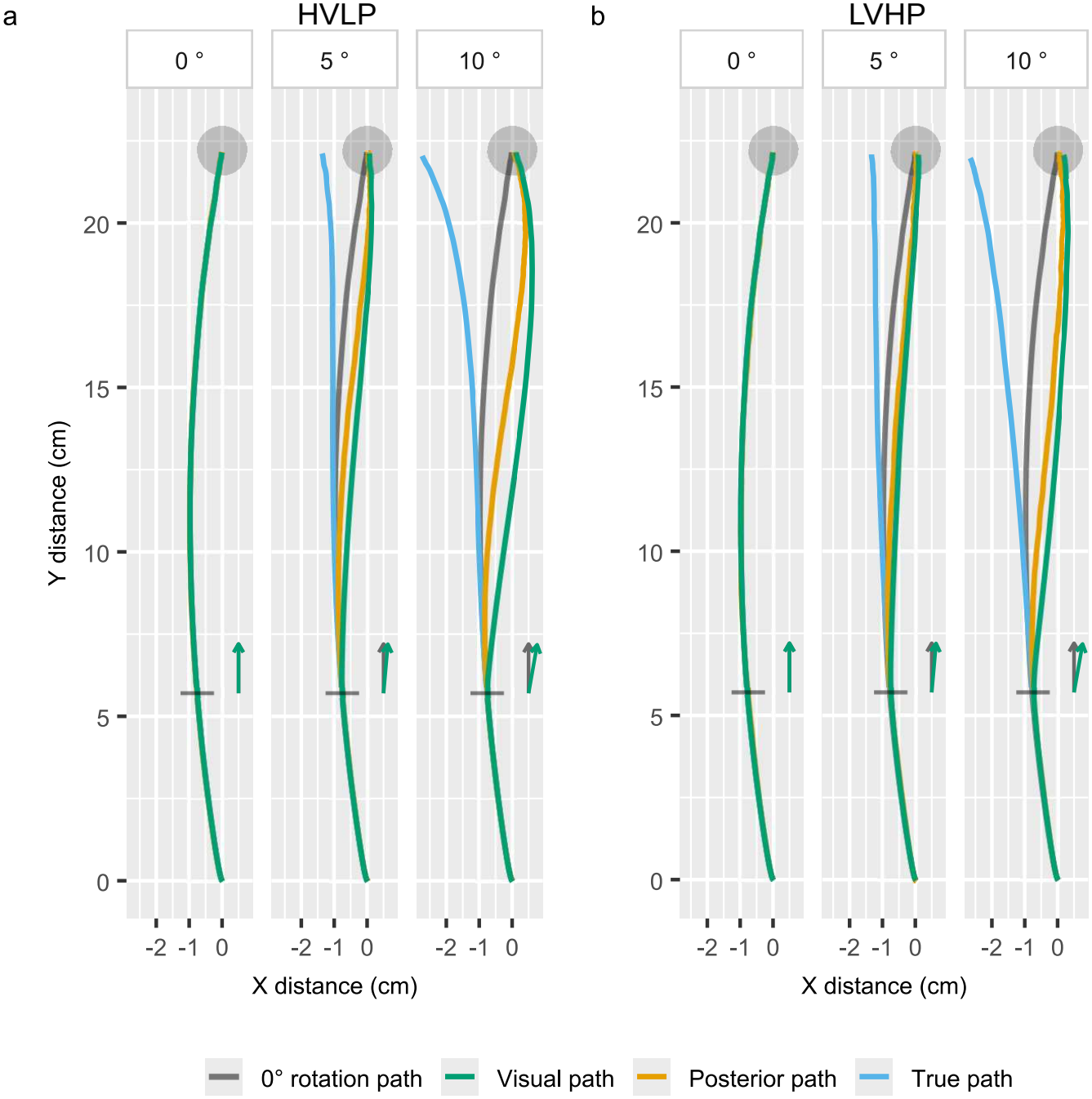
Results of simulation 2. Shows the simulated effect of applying a clockwise visual rotation of 0, 5, or 10° starting 5.7 cm into a reaching movement towards a target at a distance of 22.2 cm, mirroring the setup of Fourneret and Jeannerod (1998). Panel **a** shows the HVLP configuration, while panel **b** shows LVHP configuration. The blue trace shows the true hand path, while the posterior inferred path of the Agent is shown in orange. The true path of the unperturbed (0° rotation) condition is added in grey to facilitate comparison of how the rotated visual feedback (dark green) affects the reaching movement. The green arrows indicate the rotation applied in each plot. The paths shown here represent the averages of 500 simulated trials per condition.

The figure shows almost identical movement paths and sensory integration for both the HVLP configuration (Fig. 8**a**) and the LVHP configuration (Fig. 8**b**). Both configurations appear to match the perceptual experience reported by the majority of research participants in the study by Fourneret and Jeannerod (1998); the Agent-perceived posterior state deviates in the same direction as the visual feedback, despite the fact that both efferent motor commands and proprioceptive feedback would drive the posterior towards the true hand location, which is in the opposite direction. From the Agent’s point of view, it observes that it is moving unintendedly too far to the right as evidenced by the visual feedback, and it corrects for this “mistake” by moving slightly more in the opposite direction.

Three different sources of information influence the posterior arm state: partly forward prediction based on efferent motor commands and the previous state estimate, partly visual feed-back of hand location, and finally, proprioceptive feedback, which can be further broken down into position and velocity components. As the Agent is set up to assume the same causal origin of all three sources of information, the posterior perceived path is a weighted average of each source of information; when sensory feedback is not biased, this allows for filtering of sensory feedback and efferent output, such that the perceived arm state tracks the true arm state as well as possible, given the present sensory and motor noise. However, when incongruency is introduced through visual rotation, the posterior will then tend to be dominated by the most trusted source of information, which in this case is visual feedback. As rotation of the visual feedback is initiated after the 5.7 cm mark, the posterior state estimate of the Agent begins to show a deviation in the same direction as the offset visual feedback at this point. Reacting in response to this perceived deviation, the Agent begins to modify its efferent motor commands, so as to reach the target from its current posterior state estimate, here moving slightly more to the left than for the unperturbed trials.

Our model thus predicts that in cases such as this, where visual feedback is continuously available, it makes little difference whether proprioceptive feedback is mainly in the form of velocity-based (Fig. 8**a**) or position-based (Fig. 8**b**) information, as the visual feedback ends up dominating the posterior state estimation in either case, producing very similar outcomes.

### 4.3 Simulation 3: Reacting to brief offset visual feedback during reaching

In Fig. 9 we show the results of Simulation 3. In panels **a** and **b**, we plot out the averaged true hand path, the posterior Agent-perceived path, and the cursor path, here for an example condition where the cursor is offset 1 cm to the right. Three different visual conditions are plotted, each showing no blur, medium blur, and high blur of the visual feedback that the Agent receives. The HVLP configuration is shown in Fig. 9**a** and **c** while the LVHP configuration Fig. 9**b** and **d**.

**Fig. 9.**
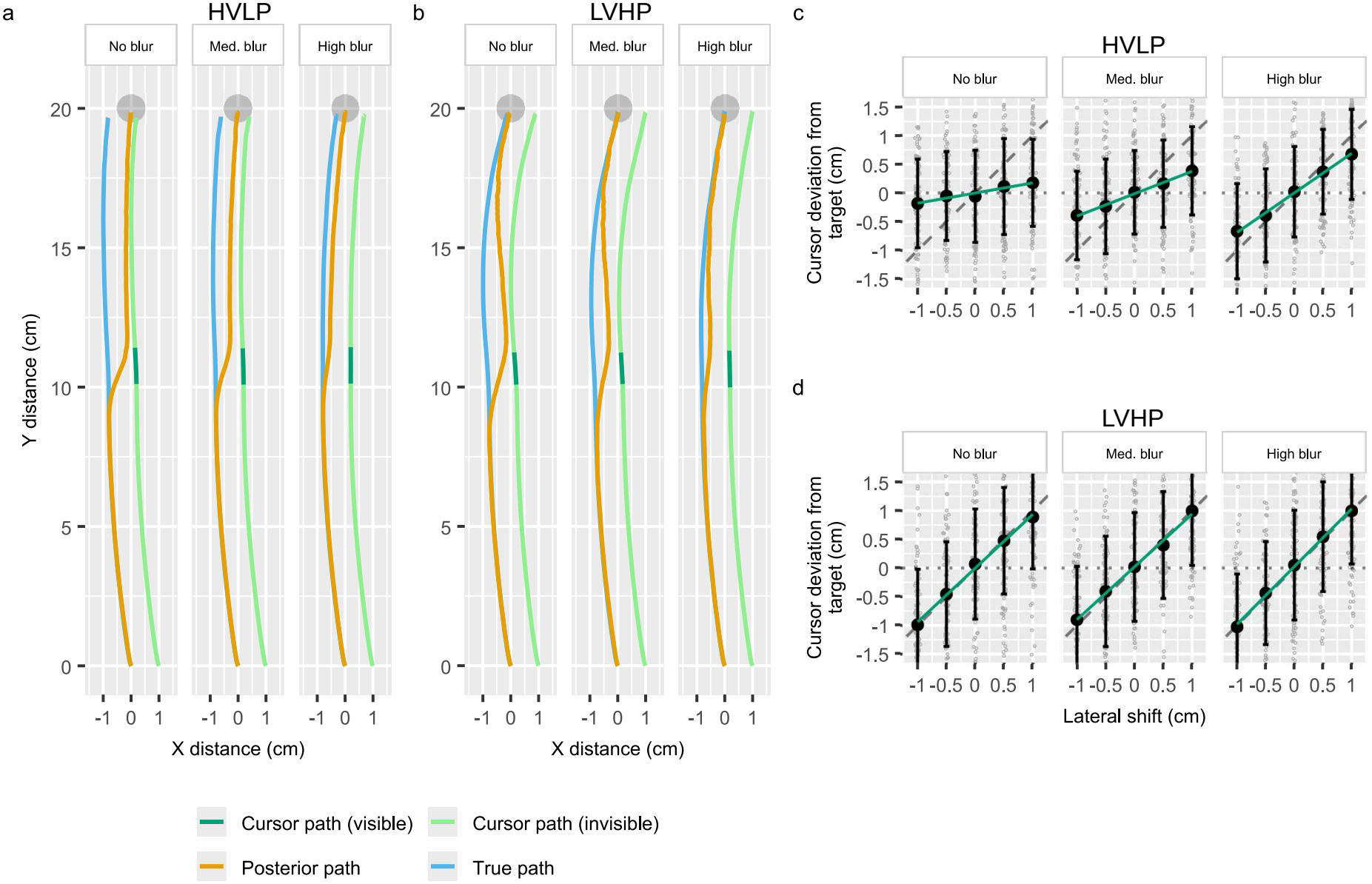
Results of simulation 3. Panels **a** and **b** shows the hand movement path for a simulated reaching task similar to the experiment performed by Körding and Wolpert (2004). Panel **a** shows the HVLP configuration, while panel **b** the LVHP configuration. The blue trace shows the true hand path, while the orange trace shows the posterior inferred path of the Agent. Both **a** and **b** display a condition where a 1 cm visual offset to the right is applied to the cursor (light green), which remains visible for only 100 ms after passing the 10 cm mark (dark green). Three sub-panels are shown, one each for no visual blur, medium visual blur, and high visual blur. Plotted paths represent the means across 500 simulated trials per condition. Panels **c** and **d** each correspond to **a** and **b** respectively, and show the endpoint cursor deviation (X distance) from the target plotted for five different lateral shifts of the cursor. Small grey dots are individual trials (subset of 100 per condition is shown), large black dots and error bars are mean +/-SD, and green lines are linear ordinary least squares regression lines. The diagonal dashed line corresponds to no endpoint correction by visual feedback, while the horizontal dotted line corresponds to full correction (no cursor deviation from the target).

For both Agent configurations, the initial 10 cm of the movement appears identical. When an offset to the position of visual feedback is applied, the forward kinematic mapping is no longer valid, resulting in a discrepancy between the predicted visual feedback and the observed values. A consistent offset in the provided visual feedback will gradually cause the Agent to update its state estimate to one that would be more likely to produce the visual observations; it may be seen how the posterior path is “pulled” towards the cursor during the brief window of visual feedback. The rate at which the posterior is pulled in this direction depends upon the relative precision of the different sources of information; this is seen clearly when medium and high visual blur are compared to the no blur condition.

After visual feedback is again removed, the simulation results show substantial differences in how the posterior state, and consequently the performed movement, is affected, depending on the configuration of the proprioceptive feedback. The HVLP Agent completes the movement with essentially the same constant offset (Fig. 9**a**), due to its reliance on high-precision velocity-based proprioceptive feedback. This configuration of proprioceptive feedback allows for high-precision estimation of changes in joint angles, while only a weak signal is available for sensing absolute joint angles. The visual-induced offset in perceived joint angles is therefore mostly kept constant over the second half of the movement. This produces a pattern of endpoint errors as a function of lateral offset and visual precision (Fig. 9**c**) that closely matches those observed by Körding and Wolpert (2004). The LVHP Agent, on the other hand, has relatively precise position sensors, while velocity feedback and, therefore, changes in joint angles, are only weakly contributing to the perceived angular positions. This has the effect of “pulling” the posterior arm state back towards the true arm state during the second half of the movement. When the target is reached, the perceptual offset induced by the visual feedback is almost washed out (Fig. 9**d**), a pattern of results that is substantially different from those reported by Körding and Wolpert (2004).

The endpoint precision of the two Agent configurations is approximately matched, with a lateral error distribution of 0.71 cm SD for the HVLP Agent and 0.77 cm SD for the LVHP Agent (for the baseline condition with no blur and no lateral offset).

## 5 Discussion

Theories based on the concepts of Bayesian sensory integration and state estimation have become central to the field of sensorimotor control over the last few decades (Todorov and Jordan 2002; Knill and Pouget 2004; Franklin and Wolpert 2011; Friston 2010). In this study, we set out to investigate the impact of two contrasting configurations of proprioceptive feedback in the context of Bayesian state estimation: one that relies primarily on velocity-based feedback (high-precision velocity, low-precision position: HVLP) and one that primarily relies on position-based feedback (lowprecision velocity, high-precision position: LVHP). Including proprioceptive sensors beyond purely position-based inputs, and specifying their precision, is an important modelling choice that is often omitted for the sake of simplicity and readability. For example, the paper by Friston et al. (2010) is one of the most cited articles on Active Inference and describes oculomotor and arm-reaching examples that use position-only proprioceptive input. While Todorov and Jordan (2002) do include velocity sensors in their simulation model, which do augment the positional estimation of their simulated Agent to some extent, the relative weighting of position and velocity-based feedback is not discussed.

In Simulation 1, we demonstrate that both the HVLP and LVHP proprioceptive feedback configurations can result in human-like adjustments in movement speed and endpoint errors when the effects of antagonist muscle vibration are simulated. Both of these configurations predict the observations made by Cody et al. (1990), showing progressively greater undershooting of a target location with increasing intensity and duration of the simulated vibration, with corresponding slowing of the movement.

These responses depended on how muscle vibration was implemented; for the HVLP configuration, vibration was implemented as a joint velocity sampling bias, whereas for the LVHP configuration, it was implemented as a joint position sampling bias. Muscle/tendon vibration acts to some degree on both type II and type Ia muscle spindle afferents, each primarily signalling position and velocity values, though with some overlap, with the effect on each driven by both vibration frequency and amplitude (Burke et al. 1976; Roll and Vedel 1982; Roll et al. 1989). Any effects observed in a given muscle vibration study must at least be considered to be partly caused by biasing of a positional and/or velocity feedback signal. While we simulate and show the individual effects of either a position or velocity offset, it seems likely that actual muscle vibration would cause both at the same time; implementing such dual-action in the two presented proprioceptive configurations would amplify the effects slightly, with the majority of the effect still mediated by the most precise of the two modes of proprioceptive feedback. Our results highlight the need for careful design and interpretation when utilising muscle vibration to investigate the sensorimotor system, as fundamentally different effect pathways are here predicted to cause very similar observed patterns of behaviour in the context of Bayesian sensory integration. Additionally, researchers using similar simulation models should carefully consider their implementation of proprioceptive feedback, as both velocity-based and position-based implementations of proprioception can produce human-like behaviour under simulated interventions such as these. While the addition of either feedback stream can augment the inference of the other, there are fundamental differences between whether information enters the inference process as rate-of-change or absolute values.

Similarly, Simulation 2 shows negligible differences in the simulated results between the HVLP and LVHP proprioceptive configurations, as visual feedback of the hand location tends to dominate the posterior state estimate in both cases. Fourneret and Jeannerod found that most participants in their study reported experiencing movement in the same direction as the offset visual feedback, rather than in the direction corresponding to the actual movement of their hand as they corrected for the offset visual feedback (Fourneret and Jeannerod 1998). For both configurations in our simulation, the posterior state estimate is updated in much the same way when visual feedback is rotated, leading to similar motor corrections. If we consider the posterior state estimate from the here-presented simulations as what might be available for conscious perception and report, then our simulations would predict the same pattern of reports. While both proprioceptive feedback and efferent output would tend to pull the posterior state estimate in the same direction as the actual movement of the hand, the high precision of visual feedback used in our simulation results in the posterior path of the hand being very close to the path of the rotated visual cursor. It has generally been found that in instances with conflicting proprioceptive and visual feedback, only a single, combined state estimate is experienced, rather than one based on visual feedback and one based on proprioceptive feedback; when available, visual feedback tends to dominate the posterior state estimation, unless its precision is intentionally reduced (Lackner and Taublieb 1984; Körding and Wolpert 2004; Chancel et al. 2016a,b; Chancel and Ehrsson 2023).

In contrast, Simulation 3 reveals fundamental differences in the behaviour predicted by velocity-dominated proprioceptive feedback compared to position-dominated proprioceptive feedback. Here we attempted to replicate the setup presented by Körding and Wolpert (2004). They elegantly demonstrated how such briefly presented visual feedback could induce an offset in the perceived position of the hand, as indicated by the effects on the endpoint lateral errors. By further manipulating the precision of the visual feedback, they illustrated how such visual feedback appears to be integrated in a Bayes-optimal fashion, with increasingly blurred visual feedback inducing smaller offsets to the perceived position of the hand. It can additionally be inferred from their results, that the induced perceptual effects are at least somewhat temporally stable; even though visual feedback is only available near the midpoint of the movement, the effects remain apparant at the conclusion of the movement.

This would tend to indicate that whatever the precision of proprioceptive positional feedback is, it is in this case insufficient to effectively correct for the induced offset over the second half of the reaching movement; likewise, we can also infer that the positional precision of unblurred visual feedback is high enough that even 100 ms is sufficient to allow for close to full correction of the movement trajectory, bringing the inferred cursor position to the target location (Körding and Wolpert 2004).

Our simulations of the HVLP proprioceptive configuration explain this pattern of relative movement through its heavy reliance on velocity-based proprioceptive feedback. Once the offset is induced through the biased visual feedback, it is effectively maintained, and the relatively low-precision positional feedback is insufficient to correct for the induced offset following removal of visual feedback. In contrast, the high-precision positional feedback of the LVHP configuration is sufficient for the induced state offset to quickly drift back towards the true state of the arm after the biasing visual feedback is removed. By the time the target is reached, the state estimate is no longer affected by the out-of-date offset visual feedback.

We further manipulated the Agent-perceived precision of the visual feedback in order to approximate the usage of blurred visual feedback employed by Körding and Wolpert. While this approach of manipulating the Agent-perceived precision of visual feedback is necessarily a simplification of the method used by Körding and Wolpert to reduce reliance on visual feedback, Limanowski and Friston (2020) have previously employed a similar simulation approach to affect the relative attention applied to proprioceptive and visual cues, which resulted in a good match to their empirical observations; we use it here to decrease the attention to visual feedback, indicating the belief that blurred visual feedback is trusted less. This simulation approach produces results that closely resemble the observations by Körding and Wolpert (2004), as well as a more recent replication (Hewitson et al. 2018). Here, our simulation further reveals that decreasing the perceived precision of the visual feedback reduces the rate of information integration from the visual source. This duration dependence of information is central to the formulation of our simulation model; being exposed to a given sensory channel for longer is here inherently more informative. Such duration dependence of visual precision has previously been indicated in terms of the localisation accuracy of visual stimuli at various durations (Adam et al. 1993, 2007; Zimmermann et al. 2013); the precision of visual stimulus localisation steadily increases over time, plateauing at around 300–500 ms. A future study investigating the interaction effects of visual blur and exposure duration during a motor task could further explore this point, as our model predicts that a longer exposure time with a given visual blur can produce the same endpoint error deviation as a lesser blur, shown for a shorter duration.

Here, it is worth noting that the endpoint deviation in this simulated task also depends on the Agent’s movement speed. The results presented here are based on simulated reaches with a duration of 2 s. While Körding and Wolpert do not report the movement duration characteristics in their study, 2 s may well be longer than the average duration in their study. Simulating the same task with a higher movement speed would tend to decrease the differences between the HVLP and LVHP configuration as they are here defined: partly, visual feedback would remain available until the hand is closer to the target, and partly, there would be less time to accumulate proprioceptive sensory evidence after visual feedback is removed and before the target is reached. As such, the LVHP configuration could also match the results reported by Körding and Wolpert quite well, given different assumptions about the utilised movement speed.

We employed what might turn out to be fairly extreme differences in the relative reliance on position- and velocity-based proprioceptive feedback, in order to highlight the different movement characteristics that would be expected of each. It seems likely that reality reflects a more even mixture, which may further depend upon the task demands. It has, for example, been indicated that it is possible to attend differentially to position- and velocity-based proprioceptive cues (Sittig et al. 1985), and similarly to focus either on visual or proprioceptive feedback (Limanowski and Friston 2020), both affecting the perceived state of the arm.

These simulation results should therefore not be taken as strong evidence in favour of the human proprioceptive system more closely matching the HVLP configuration. However, studies using a longer series of subsequent movements without visual feedback have demonstrated that while absolute proprioceptive position estimation is prone to substantial drift over time, the execution of individual relative movements continues to be well-formed, with minimal variation in movement length and direction (Brown et al. 2003a,b; Smeets et al. 2006; Patterson et al. 2017). Similarly, viewing the hand start location before a reaching movement increases the endpoint precision (Rossetti et al. 1994; Elliott and Calvert 1990; Vindras et al. 1998). Such results would tend to support the notion that relative movements are well represented by the nervous system, whereas precisely estimating absolute end-effector position in a visual reference frame appears to be a substantial challenge as the time since the last visual feedback increases.

Future experiments that combine the approach of Körding and Wolpert (2004) with long-duration movement tasks (e.g. Brown et al. 2003b; Smeets et al. 2006; Patterson et al. 2017) thus seems well positioned to further probe the relative reliance on rate-of-change versus absolute position signals. Such experiments could examine the time course of endpoint errors following brief initial offset visual feedback; quickly discarding visual offsets would tend to indicate minimal reliance on velocity-based feedback, while retaining such an induced offset for longer durations would support the notion of substantial reliance on velocity-based feedback for state inference.

### 5.1 Limitations

A key limitation of this simulation setup is that we do not include any delay of afferent feedback and efferent output; all sensory samples are generated in real time during each time step, and are immediately available for inference, and the computed torque is likewise immediately applied to the arm. This approach was chosen to limit the complexity of the model, but it has some implications which are worth considering. Much like a velocity estimate may be used to predict changes in position, an estimate of acceleration due to intended motor torque and expectation of friction is likewise useful for predicting changes to both velocity and position. This is relevant, as an extension to the model to include afferent and efferent delays would make the model considerably more reliant on its forward predictions. For example, if a 50 ms delay was added to both proprioceptive feedback and efferent motor commands, then the filter would be required to perform a 100 ms forward prediction, rather than the current single step of 10 ms, in order to bridge the temporal gap between the out-of-date sensory feedback and the point in the future when the next efferent command can affect the arm. This would affect the relative reliance on sensory feedback and efferent output for state estimation, and would necessitate different tuning of the sensory and/or motor noise parameters. Similarly, it would, for example, be very reasonable to include force-based measurements, analogous to what might be signalled by Golgi tendon organ afferents, which would add additional state-dependent information, again requiring different tuning of sensory/motor precision to achieve similar behaviour as in the current implementation. This again highlights that the specific values chosen for the simulations here are necessarily very specific to the model implementation presented here; there are many reasonable modelling choices that would require different parameter tuning. This means that while the parameters are to some extent biologically interpretable, their interpretation should be done exclusively in context. Instead, we aim to highlight how the relative reliance on position- and velocity-based proprioceptive feedback alters the pattern of inference and the resulting motor behaviour.

## 6 Conclusion

In summary, we presented an implementation of a computational model of sensory integration during active movements of a 2D planar arm, building on the principles of recursive Bayesian sensory integration. We exemplify the behaviour of the simulated Agent by comparing it with the reported results of three classical studies of sensorimotor control, demonstrating that the model can produce human-like motor behaviour under various visual and proprioceptive interventions.

With this model, we particularly highlight the importance of carefully considering the type and relative precision of the various channels of sensory feedback that a computational model is given access to, as different sets of assumptions can produce similar expected behaviour, matching human data, but requiring different interpretations. In Simulation 1, we demonstrate how different assumed effects of muscle/tendon vibration can produce very similar observed movement behaviour, matching empirical data, depending on the assumptions of the relative precision of position- and velocity-based proprioceptive feedback. This emphasises the care required when interpreting human data from such experiments. In Simulation 2, we show how the posterior arm state evolves in response to continuously displayed biased visual feedback, and discuss how the posterior of the Bayesian sensory integration process fits with the reported observations by Fourneret and Jeannerod (1998). In Simulation 3, we present how a setup using briefly presented offset visual feedback shows significant differences in the expected distribution of endpoint errors, depending on the relative weighting of position- and velocity-based proprioceptive cues. In particular, we highlight how high-precision velocity feedback allows for perception and execution of relative movements, as is seen in the study by Körding and Wolpert (2004), whose experimental setup was the basis for this simulation.

While our simulated results tend to support a significant contribution of velocity-based proprioceptive feedback for positional state inference, further research is needed to corroborate this claim.

## Statements and Declarations

### Supplementary information

Not applicable.

### Ethics approval and consent to participate

Not applicable.

### Consent for publication

All authors consent to the publication of this manuscript.

### Data availability

Simulated data sets are available at https://osf.io/kvtcu.

### Materials availability

Not applicable.

### Code availability

The simulation code is available at https://github.com/CoInAct-group/BayesianInferenceArm and archived at https://osf.io/kvtcu.

### Author contributions

ESM, MEO, and MSC all conceptualised the work. ESM and MEO both developed the modelling framework. ESM drafted the manuscript. ESM, MEO, and MSC all revised and approved the final manuscript.

## Acknowledgements

This project was supported by a DATA+ grant from University of Copenhagen. ESM, MEO and MSC were supported by the Carlsberg Foundation (CF22-0941). MSC was further supported by Independent Research Fund Denmark (DFF-FKK, 0132-00141B).

## Appendix A Agent posteriors for Simulation 1

**Fig. A1.**
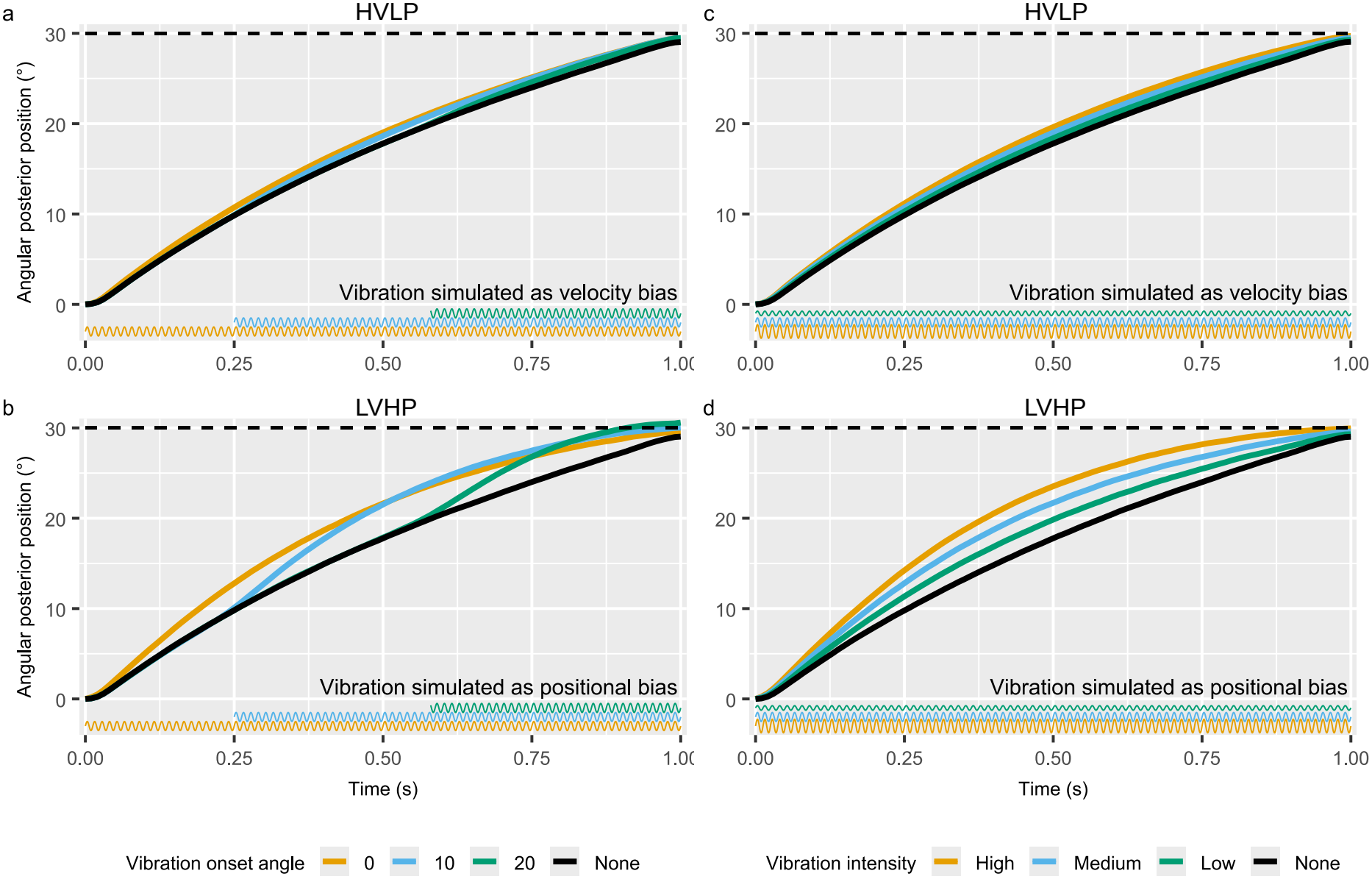
Extended results of Simulation 1. Shows the same plot as Fig. 7, but displays the Agent-perceived posterior values rather than true arm values; across all conditions, the Agent correctly guides its posterior hand to the target location, despite perturbations

## Appendix B Simulated Agent parameter values

**Table B1.**
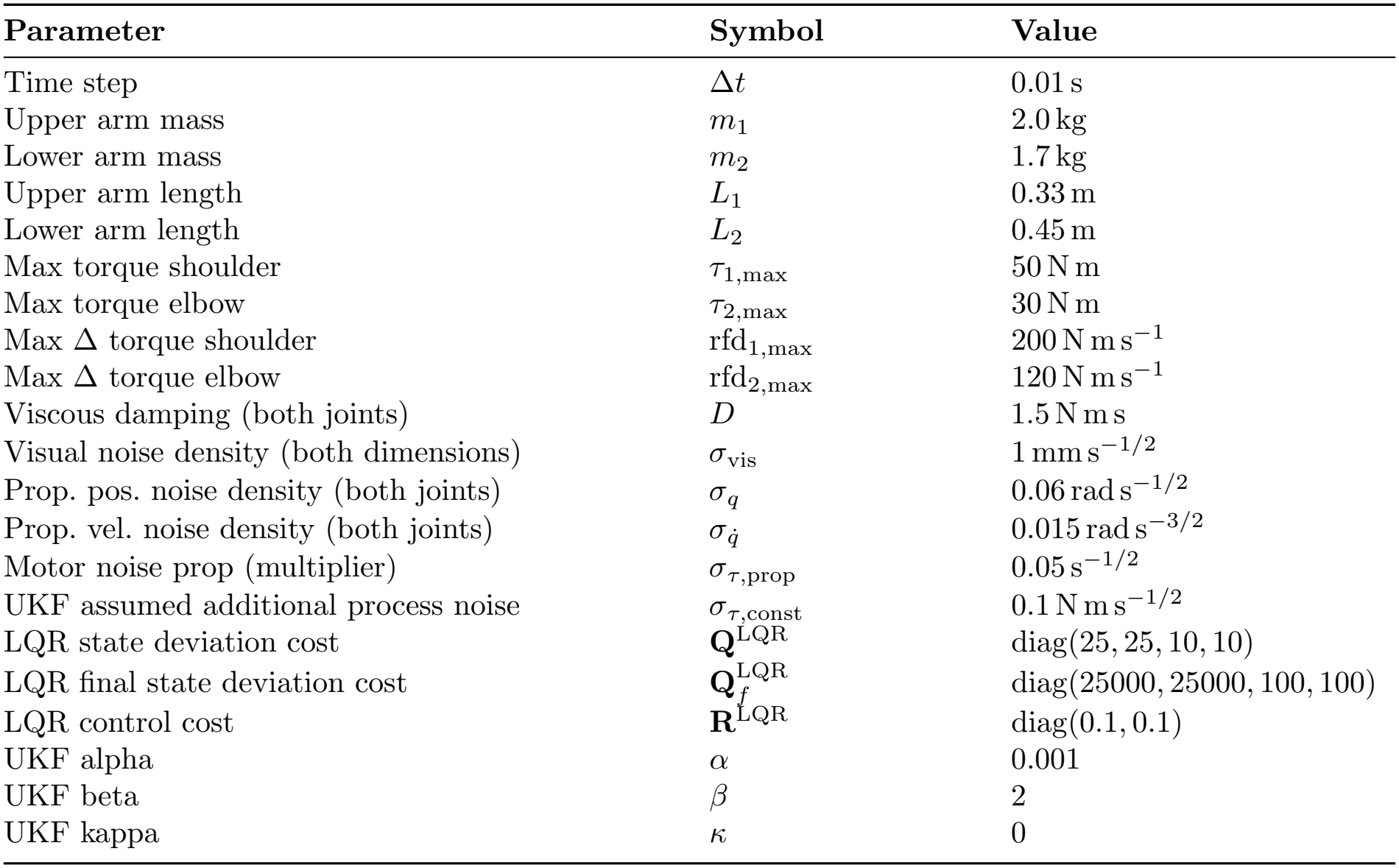
Table of simulation parameters. Noise densities are continuous-time; per-step SDs use division by 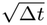. Proprioceptive pos. and vel. noise densities are here specified for the HVLP configuration; the amplitudes are flipped for the LVHP configuration.

| **Parameter** | **Symbol** | **Value** |
|---|---|---|
| Time step | Δ*t* | 0.01 s |
| Upper arm mass | *m*<sub>1</sub> | 2.0 kg |
| Lower arm mass | *m*<sub>2</sub> | 1.7 kg |
| Upper arm length | *L*<sub>1</sub> | 0.33 m |
| Lower arm length | *L*<sub>2</sub> | 0.45 m |
| Max torque shoulder | *τ*<sub>1,max</sub> | 50 N m |
| Max torque elbow | *τ*<sub>2,max</sub> | 30 N m |
| Max Δ torque shoulder | rfd<sub>1,max</sub> | 200 N m s<sup>−1</sup> |
| Max Δ torque elbow | rfd<sub>2,max</sub> | 120 N m s<sup>−1</sup> |
| Viscous damping (both joints) | *D* | 1.5 N m s |
| Visual noise density (both dimensions) | *σ*<sub>vis</sub> | 1 mm s<sup>−1/2</sup> |
| Prop. pos. noise density (both joints) | *σ*<sub>*q*</sub> | 0.06 rad s<sup>−1/2</sup> |
| Prop. vel. noise density (both joints) | *σ*<sub>*q̇*</sub> | 0.015 rad s<sup>−3/2</sup> |
| Motor noise prop (multiplier) | *σ*<sub>*τ*,prop</sub> | 0.05 s<sup>−1/2</sup> |
| UKF assumed additional process noise | *σ*<sub>*τ*,const</sub> | 0.1 N m s<sup>−1/2</sup> |
| LQR state deviation cost | **Q**<sup>LQR</sup> | diag(25, 25, 10, 10) |
| LQR final state deviation cost | $\mathbf{Q}_f^{\mathrm{LQR}}$ | diag(25000, 25000, 100, 100) |
| LQR control cost | **R**<sup>LQR</sup> | diag(0.1, 0.1) |
| UKF alpha | *α* | 0.001 |
| UKF beta | *β* | 2 |
| UKF kappa | *κ* | 0 |

